# Resting State Networks in empirical and simulated dynamic functional connectivity

**DOI:** 10.1101/089516

**Authors:** Katharina Glomb, Adrián Ponce-Alvarez, Matthieu Gilson, Petra Ritter, Gustavo Deco

## Abstract

It is well-established that patterns of functional connectivity (FC) - measures of correlated activity between pairs of voxels or regions observed in the human brain using neuroimaging - are robustly expressed in spontaneous activity during rest. These patterns are not static, but exhibit complex spatio-temporal dynamics. Over the last years, a multitude of methods have been proposed to reveal these dynamics on the level of the whole brain. One finding is that the brain transitions through different FC configurations over time, and substantial effort has been put into characterizing these configurations. However, the dynamics governing these transitions are more elusive, specifically, the contribution of stationary vs. non-stationary dynamics is an active field of inquiry. In this study, we use a whole-brain approach, considering FC dynamics between 66 ROIs covering the entire cortex. We combine an innovative dimensionality reduction technique, tensor decomposition, with a mean field model which possesses stationary dynamics. It has been shown to explain resting state FC averaged over time and multiple subjects, however, this average FC summarizes the spatial distribution of correlations while hiding their temporal dynamics. First, we apply tensor decomposition to resting state scans from 24 healthy controls in order to characterize spatio-temporal dynamics present in the data. We simultaneously utilize temporal and spatial information by creating tensors that are subsequently decomposed into sets of brain regions (“communities”) that share similar temporal dynamics, and their associated time courses. The tensors contain pairwise FC computed inside of overlapping sliding windows. Communities are discovered by clustering features pooled from all subjects, thereby ensuring that they generalize. We find that, on the group level, the data give rise to four distinct communities that resemble known resting state networks (RSNs): default mode network, visual network, control networks, and somatomotor network. Second, we simulate data with our stationary mean field model whose nodes are connected according to results from DTI and fiber tracking. In this model, all spatio-temporal structure is due to noisy fluctuations around the average FC. We analyze the simulated data in the same way as the empirical data in order to determine whether stationary dynamics can explain the emergence of distinct FC patterns (RSNs) which have their own time courses. We find that this is the case for all four networks using the spatio-temporal information revealed by tensor decomposition if nodes in the simulation are connected according to model-based effective connectivity. Furthermore, we find that these results require only a small part of the FC values, namely the highest values that occur across time and ROI pair. Our findings show that stationary dynamics can account for the emergence of RSNs. We provide an innovative method that does not make strong assumptions about the underlying data and is generally applicable to resting state or task data from different subject populations.

## 1 Introduction

The question of how large-scale cortical function arises from underlying anatomical connectivity has been the object of much investigation since the advent of non-invasive imaging techniques (Vincent et al., 2007; Matsui et al., 2011; Wang et al., 2013), in particular since it was discovered that interareal functional relationships found under task conditions are maintained during rest (Biswal et al., 1995; Cordes et al., 2000; Beckmann and Smith, 2004; Fox et al., 2005). With magnetic resonance imaging (MRI) it is possible to obtain both functional and structural connectivities (FC and SC, respectively). Although there is large variability across subjects and sessions, both in SC (Heiervang et al., 2006) and FC measures (Mueller et al., 2013; Finn et al., 2015), studies using group averages have revealed general principles of information processing in the brain (Raichle et al., 2001; Doucet et al., 2011; Van den Heuvel and Sporns, 2011; Deco and Jirsa, 2012; Haimovici et al., 2013).

In order to connect SC and FC, computational models are an important tool for understanding how activity propagates from one node to another to produce the observed data (Honey et al., 2009; Cabral et al., 2012; Deco et al., 2014). Most models optimize their parameters by fitting the average FC. Only recently, the question of whether and how relevant information can be extracted from the spontaneous fluctuations in pairwise FC strength, and how to describe the richness of the temporal dynamics, has received increasing attention in data analysis (Chang and Glover, 2010; Hutchison et al., 2012; Allen et al., 2012; Tagliazucchi et al., 2012; Liu et al., 2013; Leonardi and Van de Ville, 2013a; Zalesky et al., 2014; Yaesoubi et al., 2015) and modelling (Hansen et al., 2014; Ponce-Alvarez et al., 2015). This has lead to the notion of dynamic functional connectivity (dFC); dFC has been shown to be relevant for behavior (Kucyi et al., 2013; Kucyi and Davis, 2014; Barttfeld et al., 2015; Chen et al., 2015; Yang et al., 2014), development (Madhyastha and Grabowski, 2014; Hutchison and Morton, 2015; Tagliazucchi et al., 2016; Marusak et al., 2017), and disorders (Damaraju et al., 2014; Wee et al., 2016; Miller et al., 2016; Rashid et al., 2014; Sourty et al., 2016; Demirtaş et al., 2016) and is therefore likely to have a basis in neural activity.

Here, we use a dynamic mean field model (Wong and Wang, 2006) of the human cortex which has been shown to reproduce average resting state (RS) FC (Deco et al., 2014). It is our goal to determine whether simulated data exhibit FC patterns over time that resemble those of empirical data. Specifically, we want to test whether resting state networks (RSNs) can be explained in this way. To this end, we analyze RS data from 24 healthy subjects (Schirner et al., 2015) and compare to simulated data. The cortex is modelled by 66 nodes corresponding to 66 brain areas also used to parcellate the empirical data. The nodes are connected according to empirical SC derived from the same subjects via diffusion weighted MRI and fiber tracking.

We use tensor decomposition for extracting relevant and general features of the spatio-temporal dynamics. This method has been shown to work well for community detection (Gauvin et al., 2014) and has been applied to brain data (Cichocki, 2013; Leonardi and Van de Ville, 2013a; Ponce-Alvarez et al., 2015; Leonardi and Van De Ville, 2013b). Unlike ICA, which has become the standard method for extracting RSNs (McKeown et al., 1998; Beckmann et al., 2005; Mantini et al., 2007), tensor factorization does not assume spatial independence of the underlying components, which is a strong constraint not directly motivated by the data. Here, such a constraint is not required and the space of possible solutions is not unnecessarily restricted. Furthermore, it has the advantage that it can readily be used at our level of spatial resolution.

The modelling approach aims at linking FC and SC. One conceptual problem of SC is that it provides neither directionality information nor the weights of the connections. These two points are addressed by the concept of effective connectivity (EC). SC can be viewed as an approximation to EC, and it is the latter that is genuinely related to the dynamics in network models (Friston, 1994). Reversely, underlying connectivity (SC or EC) can be inferred from FC, or more generally, from the dynamics found in the data, through the same kinds of models. Gilson et al. (2016) developed a method to extract EC from RS fMRI data using a noise diffusion model which possesses simpler dynamics than the DMF. They show that the EC that accounts best for empirical FC significantly differs from the SC in a number of points. We use both SC and EC as underlying connectivity in our model and explore how their properties are linked to the spatio-temporal patterns found in empirical and simulated data.

## 2 Methods

### 2.1 Empirical data

RS fMRI as well as corresponding diffusion weighted (dw) MRI data were collected from 24 healthy participants (11 female) at the Charité Berlin, Germany, by Petra Ritter and co-workers. The original dataset consisted of 49 subjects, but we chose only those aged 18 to 35 years (mean 25.7 years) since it is known that FC changes with age (Meunier et al., 2009). Each fMRI dataset amounts to 661 time points recorded at TR=2s, i.e. about 22 minutes. In the same session, EEG was also recorded, but we do not use the data here. RS BOLD was recorded while subjects were asked to stay awake with their eyes closed, using a 3T Siemens Trim Trio scanner and a 12 channel Siemens head coil (voxel size 3 *×* 3 *×* 3mm). Voxel time courses are averaged inside ROIs defined by the Desikan-Killiany atlas (Desikan et al., 2006) as implemented in FreeSurfer. We removed the areas labeled as corpus callosum on both sides since they only contain white matter, amounting to 33 cortical ROIs for each hemisphere. See table S1 for details.

The diffusion tensors (TR=750 ms, voxel size 2.3 *×* 2.3 *×* 2.3mm) computed from the dwMRI data recorded with 64 gradient directions were subjected to probabilistic fiber tracking as implemented in MRTrix (Tournier et al., 2004, 2007) in order to obtain structural connectivity (SC) matrices for each subject. Masks derived from high-resolution T1-images were used to determine seed-and stop-locations for fibers in the grey matter/white matter-interface (GWI). SC matrices contain connection strengths which are estimated by combining the number of streamlines obtained from the fiber tracking algorithm with various assumptions based on known limitations imposed by anatomy, notably the size of the GWI of each region. We use the average over all 24 subject in our simulations.

Further details are available in Schirner et al. (2015).

### 2.2 Model data

A dynamic mean field approximation of a network of populations of leaky integrate and fire neurons was used to simulate RS activity (Brunel and Wang, 2001; Wong and Wang, 2006; Deco and Jirsa, 2012; Deco et al., 2014), as illustrated in figure 1. The excitatory populations are connected using a) an average over the SC matrices derived from dwMRI data (section 2.1), b) a heuristically enhanced version of SC with added homotopic connections, or c) an effective connectivity (EC) matrix derived from these SC matrices as described in section 2.9.

**Figure 1:**
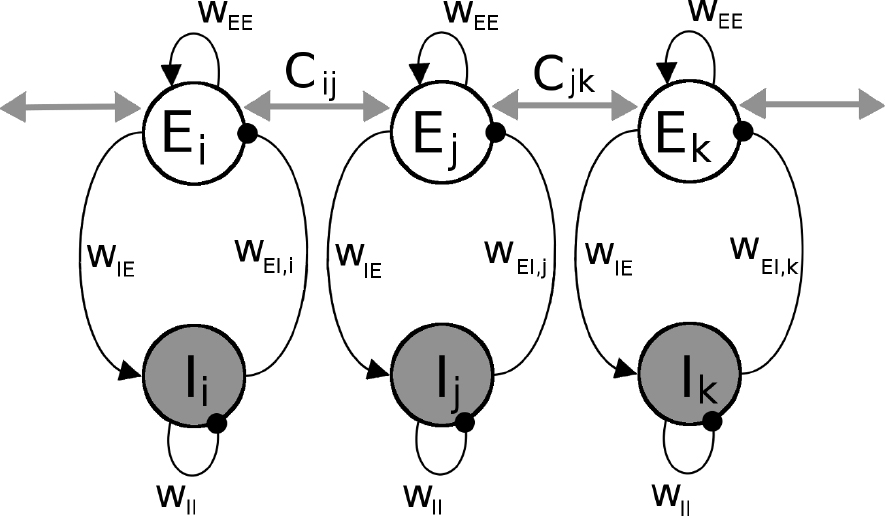
Schematic view of dynamic mean field (DMF) model used to simulate synaptic activity. Each brain area is modelled by a pair of excitatory (E) and inhibitory (I) pools. The local connectivity is governed by the four weights *w*_EE_, *w*_EI_*_,i_, w*_IE_, and *w*_II_, whereby *w*_EI_*_,i_* is adjusted for each population individually so as to keep the firing rates at a low level (3-10 Hz). Black lines with spheres signify GABA connections, black arrows, NMDA. Gray arrows are long range connections mediated by AMPA synapses and whose weights are set by the entries *ij* of the SC or EC matrix, *C*.

The activity of the populations is computed using a set of coupled nonlinear stochastic differential equations:

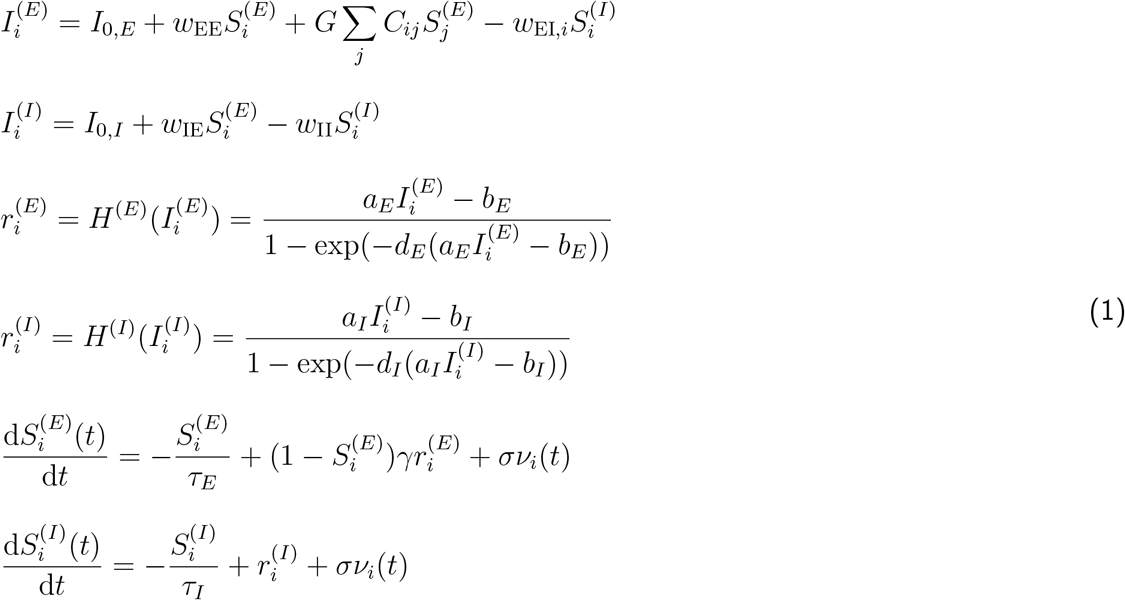

Constants are listed in table 1. Super-/subscripts *E* and *I* denote the excitatory and inhibitory pools of population *i*, respectively. *I*_*i*_ denotes synaptic input currents, which are turned into population firing rates *r*_*i*_ via sigmoid transfer functions *H*(*·*). These transfer functions and their parameters approximate the f-I curves of the full spiking model (Abbott and Chance, 2005; Deco et al., 2014) which in turn are based on experimental measurements. *S*_*i*_ denotes the average synaptic gating variable, or activity, and *ν*(*t*) is Gaussian noise with amplitude *σ* = 0.01. The kinetic parameter *γ* = 0.641 is related to the NMDA conductances’ dependence on the extracellular Mg^2+^ concentration. The time constants *τ*_*E*_ and *τ*_*I*_ are the same as in the spiking model and are based on experimental results for the rise and decay time constants of NMDA and GABA synapses (Brunel and Wang, 2001; Deco et al., 2014).

**Table 1:**
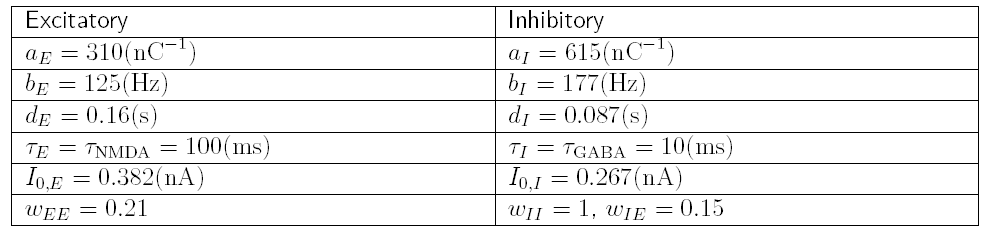
Parameters of the DMF model.

Synaptic currents *I*_*i*_ are a result of of inputs from the local network, i.e. 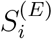 and 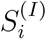, and inputs from other network nodes *j*, i.e. 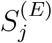. Local inputs are governed by four weights, *w*_EE_, *w*_EI_,*_i_*, *w*_IE_, and *w*_II_. Additionally, there are constant inputs to each population, denoted by *I*_0*,E*_ and *I*_0*,I*_ and corresponding to 800 Poissonian input spike trains with an average 3 Hz firing rate, weighted differently for the excitatory and inhibitory populations. Inputs from other parts of the network are provided by the excitatory populations and weighted by the entries *C*_*ij*_ for the connectivity from region *i* to region *j*, noted in the SC or EC. The diagonal of **C** is set to 0. Weights are scaled by the global coupling parameter *G*.

All local parameters are chosen such that the firing rate of a single disconnected population is close to 3 Hz (Deco et al., 2014). The one exception is the feedback inhibition, *w*_EI*,i*_, which is adjusted before each simulation using the fully connected model. This is necessary in order to ensure that the network is in its asynchronous state where we only have one stable fixed point with firing rates between 3 and 10 Hz for all regions (Deco and Jirsa, 2012). We can determine the stability of the system by taking advantage of a semi-analytical solution (Deco et al., 2014). We calculate the Jacobian matrix and confirm that all eigenvalues are negative with zero imaginary parts. Simulations were only performed for values of *G* that warranted stability of the system.

Number and length of simulations are matched to the empirical data. BOLD time courses are obtained from the synaptic activities of the excitatory pools via the Balloon-Windkessel model (Friston et al., 2003; Deco et al., 2013). Time courses are downsampled to match the TR of the experimental data.

### 2.3 Tensorization of the data

In order to take advantage of the temporal information, we adopt the widely used approach of sliding windows to compute time-dependent dynamic FC (dFC). We use overlapping windows *w* of width 120s (60 data points, TR=2s) that we advance along the time course in increments of 2s (1 frame), which results in *W* = *T −* 60 windows for each dataset (subject or simulation), *T* being the number of frames. For each window, we compute an *N × N* matrix of pairwise connectivity values, *dFC*(*w*). By concatenating these matrices along the temporal dimension, we represent each dataset as a tensor of dimensions *N × N ×W* (see figures 2A and B).

**Figure 2:**
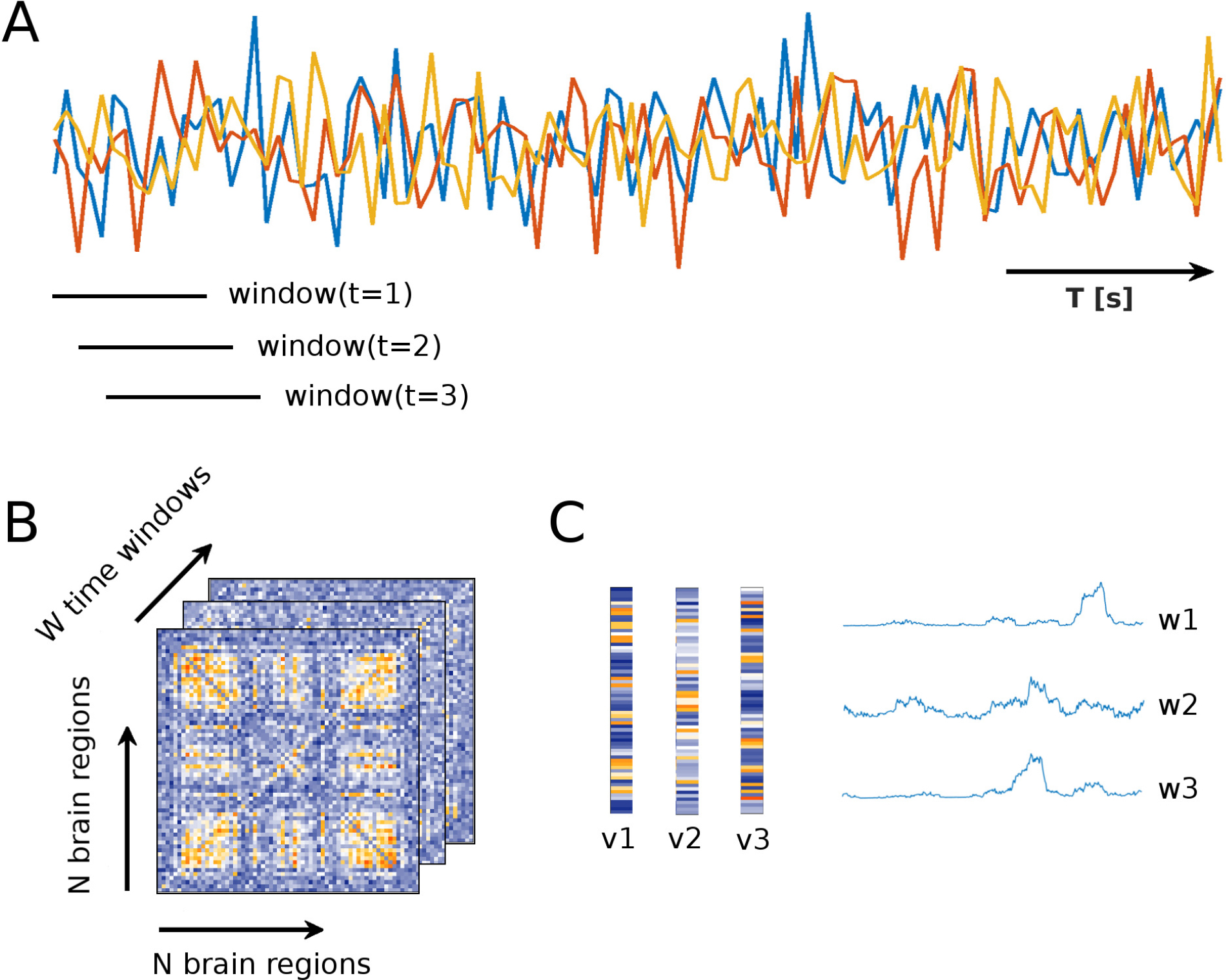
Overview of tensor decomposition. **A** We use overlapping sliding windows of width 120s (60 frames) starting at time points *t* = 1,2, …,W. They are moved along the BOLD time courses (shown schematically for three ROIs). **B** We compute time dependent dFC(w)-matrices inside these time windows. Each dFC(w)-matrix has dimensions *N×N, N* being the total number of brain regions. This results in an *N ×N×W* tensor. The entry at *dFCij* (*w*) is the functional connectivity value of ROIs *i* and *j* in window *t*. **C** Each tensor is decomposed into *F* “features” - the figure illustrates the case of *F* = 3. Each feature *i* consists of one vector for each dimension. Since dFC(w)-matrices are symmetric, the first two vectors, *v*^*i*^, corresponding to the spatial dimensions, are identical and of length *N*. The third vector, *wi*, is a time course of length *W*.

We use two measures of FC: on the one hand the most widely used one, i.e. Pearson correlation, on the other, mutual information (MI), a non-negative and nonlinear measure that allows us to constrain the tensor decomposition. The calculation of MI follows Kraskov et al. (2004) and is based on nearest neighbor distances, thus being adaptive and continuous.

In short, when estimating the MI between the 60 data points inside our window, we determine he distance between any two points [*x*_*i*_*, y*_*i*_] and [*x*_*j*_*, y*_*j*_] with *i*≠*j* and take the maximum norm, i.e.

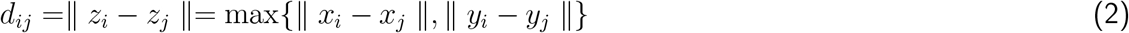

The nearest neighbor to each point [*x*_*i*_*, y*_*i*_] is the point with the minimum *d*_*ij*_, and we term this distance *ɛ*_*i*_. For each point [*x*_*i*_*, y*_*i*_], we count how many points are within this minimum distance *ɛ*_*i*_, separately for the x- and y-directions, resulting in two numbers *n_x_*(*i*) and *n_y_*(*i*). We estimate MI as 
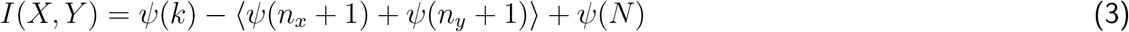
where *X* and *Y* are the two time series, and *ψ*(*·*) denotes the digamma function 
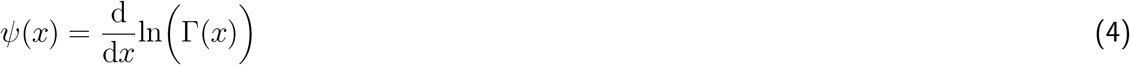
*k* = 1 because we consider the nearest neighbor; *N* is the number of data points, i.e. 60. Since this is a continuous measure, it is possible for *I*(*·, ·*) to become negative whenever the following inequality is satisfied:

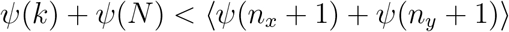

This happens when there is very little MI and many points are closer than the nearest neighbor. In these cases, we simply set MI to zero.

### 2.4 Extracting communities from low resolution fMRI data

We apply tensor factorization to both the empirical and the model data (figure 2C). The problem for the three-dimensional case treated here can be formulated as follows (Cichocki et al., 2009):

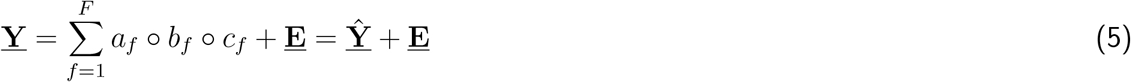
where **<u>Y</u>** is the data tensor of dimensions *N×N×W*, defined as in section 2.3. *F* is the number of features we wish to extract. *o* denotes the outer product. **A** = [*a*_1_*, a*_2_*, …, a*_*f*_], **B** = [*b*_1_*, b*_2_*, …, b*_*f*_] and **C** = [*c*_1_*, c*_2_*, …, c*_*f*_] are the factor matrices that contain the features of each of the three dimensions, respectively. Here, **A** = **B** due to the symmetry of the dFC matrices. They contain *F* vectors of length *N* with weights that are interpreted as membership values for a community. The third set **C** contains their associated time courses and is of dimensions *F×W*.

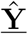 is an approximation of the data based on the features, and **<u>E</u>** is the error/noise not described by the features. Hence, the distance between **<u>Y</u>** and 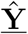 can be used to assess how well the extracted features approximate the data. We use the Frobenius norm in the case of continuous, non-thresholded tensors and the Hamming distance for thresholded, binarized tensors (section 2.5). Note that in the latter case, the result of the decomposition is continuous although the input is binary, so we threshold and binarize the reconstructed tensor such that the number of ones is preserved.

This technique is based on the very general Tucker model which can be viewed as a generalization of SVD to tensors. Unlike for SVD, though, convergence to a unique and optimal solution is not guaranteed. Consequently, it is impossible to determine the true rank of the tensors and thus, the appropriate number of features *F*. This problem is mitigated by the inclusion of further constraints, in this case, non-negativity when using absolute value of correlation or MI to construct the tensors (section 2.3). Furthermore, we use clustering of the obtained spatial features in order to ensure that they generalize across subjects and thus describe true properties of the data (section 2.7).

To decompose tensors constructed using correlation, we use the algorithm described in Phan et al. (2013). For the non-negative measures (absolute value of correlation, mutual information), we apply non-negativity of the resulting features as an additional constraint and use the algorithm described in Kim and Park (2012). Both algorithms are implemented in Matlab, requiring the tensor toolbox (Bader et al., 2015; Acar et al., 2011), and available on-line.

### 2.5 Thresholding

We reduce noise by thresholding and binarizing the tensors. This approach was also chosen in Ponce-Alvarez et al. (2015). It would perhaps be most desirable to use only the dFC values that are significant. However, it is too time demanding to generate the appropriate number of surrogate datasets to achieve the desired significance level to account for the high number of ROI pairs, windows and subjects/simulations. Hence, we simply use the x-th percentile as a threshold *θ*, where *x* = *{*0, 75, 80, 90, 91*,…*, 99*}* and, for *x >* 0, set all elements *Y*_*ijt*_ of tensor **<u>Y</u>** which are bigger than or equal to that percentile to one and everything else to zero. Note that we do not make any claims about non-stationarity of the FC time courses.

### 2.6 Surrogate data

To further validate our results, we conduct analyses with surrogate data alongside those for real data. Surrogate data are constructed by removing the pairwise correlation structure of the original time series while keeping the Fourier spectrum constant. More specifically, we Fourier transform our original time courses *x*_*i*_(*t*) of region *i* for time points *t* = 1, 2*,…, T* using Fast Fourier Transform and add random phases *φ*_*r*_ to each frequency bin before transforming back.

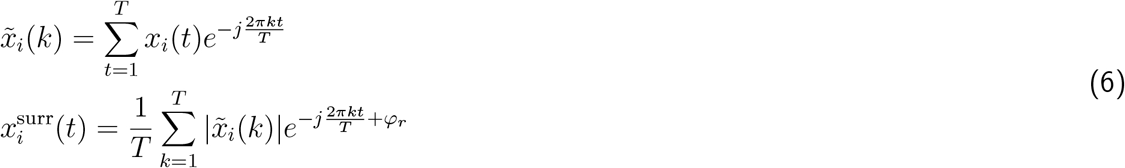
*k* = 1, 2,…*T* and *φ*_*r*_ is uniformly distributed between *−π* and *π*. This results in time courses that have the same spectral properties and autocorrelations as the original data, but are uncorrelated.

### 2.7 Constructing templates from empirical data

We construct spatial templates directly from the empirical data. First, we decompose each subject’s data tensor as described above, using a range of numbers *F* of features. In our case, the goal is to extract spatial maps that are common across subjects, so we apply a simple clustering algorithm (K-means clustering) to the set of all *F · S* features (i.e. pooled from all subjects) with different numbers *K* of clusters. We test the quality of the clustering by evaluating the mean silhouette value (de Amorim and Hennig, 2015). The resulting cluster centers of the parameter combination with the maximum silhouette value are used as templates.

This approach has two major benefits. First, finding cluster structure on the group level speaks to the robustness of our approach and validates the single subject results. This is important in light of the aforementioned problems with uniqueness and optimality of the solution which makes the true value of *F* impossible to determine. Showing that the results are similar across subjects for a certain set of parameters means that we are extracting true data structure that is not driven by individual differences. Second, our method does not have a bias towards yielding RSNs as would be the case if we were to use templates from the literature obtained with a different method, e.g. ICA on the voxel level. Therefore, showing that the templates resemble RSNs known from the literature further validates our method.

For each point *i* (here, an *N*-dimensional feature) the average distance (in terms of correlation) to points assigned to the same cluster is evaluated, denoted by *a*_*i*_, as well as the smallest average distance to all points assigned to a different cluster (i.e., the closest cluster), denoted by *b*_*i*_. Then the silhouette value is calculated as

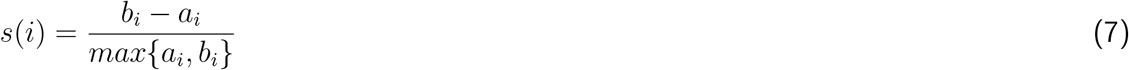

Obviously, *s*(*i*) will lie between -1 if the point is entirely in the wrong cluster, and 1 if the assignment is perfect. Taking the mean over all features gives an estimate of how well the data points are clustered.

### 2.8 Calculating the overlap between templates and features

In order to determine how well, overall, communities extracted from simulated data match those from empirical data, we compute the average overlap between all simulated spatial features with any of the templates, comparing connectivities (EC vs. SC, plus SC_e_) and using the full range of global coupling parameter values (*G*), i.e. *G* = 0.5, 0.6,…4 for simulations with the SC and *G* = 1, 1.1,…6 for simulations with the EC. For SC_e_, we use a limited range of *G*, since we are tuning an additional parameter, i.e. the weight of homotopic connections (see section 2.9).

Due to the high thresholds, the features are somewhat sparse. Therefore, correlation is not a good choice to measure the distance between features and templates. Instead, we use confusion matrices and Cohen’s Kappa. Briefly, we quantize the features on three levels and compute the overlap between any two vectors (one template, one feature extracted from simulated data) by determining the overlap for each level, creating a confusion matrix. Cohen’s Kappa is a summary of the confusion matrix:

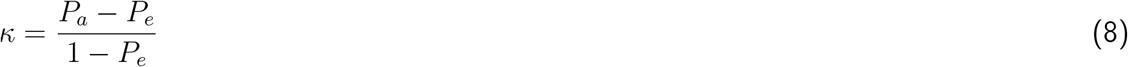
*P_a_* is the overlap, *P_e_* is the expected overlap.

As the overall match between a feature extracted from simulated data and the templates of the empirical data, we consider the mean over the maximum *κ* values for each feature in each simulation. In other words, we assume that each simulated feature corresponds to only one template. We compute an overall match for each value of *G* by averaging this value over the features and simulations.

### 2.9 Different types of connectivities

In this study, we compare simulation results from three different underlying connectivities. First, we have dwMRI-derived SC containing estimates of fiber densities from the same 24 subjects whose BOLD signals are analyzed. Second, we consider a simple enhancement of this matrix, following Messé et al. (2014), referred to as SC_e_. One of the flaws in fiber tracking is that tracts through the corpus callosum are typically underestimated or missing, resulting in strong homotopic connections being omitted. In order to directly test which effect this has, we simply add these connections in SC_e_, trying different values.Third, we use model-based effective connectivity (EC). Here, we only tune connections that are present in the SC, plus homotopic connections, and which are above a certain threshold chosen heuristically such that the level of connectivity is at 32% (Gilson et al., 2016). For SC_e_, the weights *D* of these homotopic connections are set to a unit value. We use 50 equally spaced values, the minimum being the minimum that is present for these connections in the thresholded SC, and the maximum being the overall maximum. Thus, we would in principle need to search both the range of *G* and of *D*. Simulating *S* = 24 data sets and tensorizing the resulting data takes about 48 hours; we did not have the computational resources to obtain these results. Therefore, we determine, using only one round of simulation, the range of *G* with the best fit to the average FC and then run the full 24 rounds of simulations for 5 values around this best *G*. We then determine the average overlap for all values of *D*.

In the following, we briefly describe the method developed by Gilson et al. (2016) for constructing EC matrices by combining SC and FC. The key is to extract information about cortical interactions from BOLD covariances with non-zero time shifts:

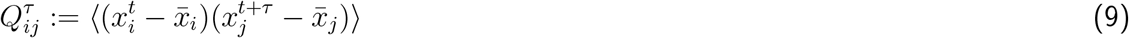

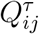 is the covariance between BOLD time courses *x* of ROIs *i* and *j*, with *x*_*j*_ shifted by *τ* against *x*_*i*_. The angular brackets denote averaging over randomness due to noise in the model, such that the mean BOLD for node *i* is 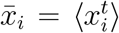. For *τ* ≠ 0, this matrix is non-symmetric. The goal is to estimate the underlying connectivity such that the model minimizes the error between model covariances (*Q*^0^ and *Q^τ^*) and their empirical counterparts (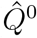 and 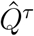), for a given *τ* equal to 1 TR.

EC is model based, meaning that there is an underlying assumption of how activity propagates through the brain using the present connections to activate the nodes. We use a noise diffusion model with a static nonlinearity, Φ:

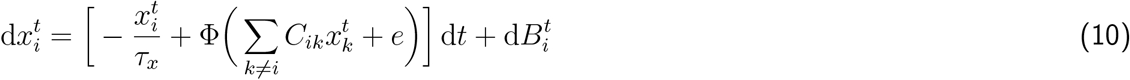

Time course *x*_*i*_ of region *i* is subject to an exponential decay with time constant *τ*_*x*_ at each time point *t*. *C* is the connectivity matrix that contains weights linking regions *i* and *j*, and the sum is over all regions *k* from which *i* receives input. This means that activation is only provided by the input from other nodes, weights for which are defined in *C*. The background input *e* is shared by all *i*. Fluctuations of the activities are driven by Gaussian noise 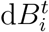. The model directly simulates BOLD activity, hence the Jacobian of the system is simply:

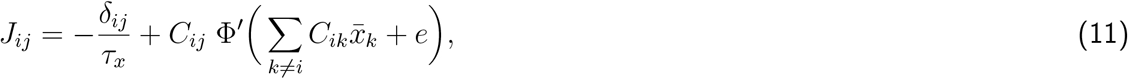
where *δ*_*ij*_ is the Kronecker delta and *Phi′* denotes the first time derivative of Φ. Therefore, the model is solely constrained by *C*.

We want to estimate *J* and therefore *C* such that it satisfies the steady state of the second order fluctuations:

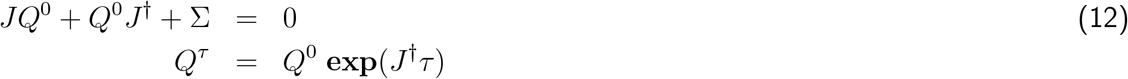

Σ is the noise matrix with diagonal terms 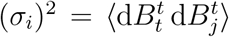; † denotes the matrix transpose and **exp** the matrix exponential. Since we are using empirical covariances estimated from fMRI data, the objective covariances 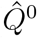 and 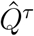 are very noisy and make a direct estimation via an analytical approach unfeasible. Therefore, we use the iterative Lyapunov optimization (LO) procedure described in Gilson et al. (2016).

The update works by simulating the BOLD activity using the model defined in equation 10 without noise and the current connectivity *C*, so as to evaluate the mean activity 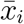 for all regions, yielding the Jacobian *J* in equation 11. Then, the model covariances *Q*^0^ and *Q^τ^* are given by equation 12, using the Bartels-Stewart algorithm for the first line and using the current values for *J* and Σ. The model covariance matrices are compared to the objective covariances 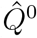 and 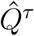. Then, the Jacobian update is evaluated according to:

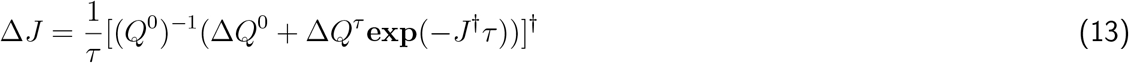

Finally, we obtain the connectivity update 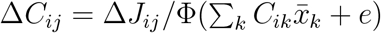.

In addition, the noise matrix Σ (see equation 12) is optimized at the same time as *C*. We assume that each node receives independent noise, meaning that Σ is diagonal. The update is performed such that the model variances coincide with the empirical values:

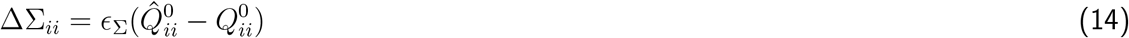

It was shown that a time shift *τ* equal to 1 or 2 TR gives a good estimation performance (Gilson t al., 2016). In fact, *τ* has to roughly match the time scale on which the neural activity decays, i.e. _*x*_ in equation 11. The latter is estimated from the slope of the autocorrelation of each region (the slope is close to 1*/τ*_*x*_), and results in *τ* = 5.3s which leads us to set the time shift to 1 TR=2s.

### 2.10 Analysis pipeline

To summarize the methodology of the paper, we give a brief overview of the steps involved in obtaining the results described in the next section. We have *S* = 24 sets of empirical resting state data (from 24 subjects) of length *T* = 661 frames (TR=2s). Voxel time courses are averaged inside of *N* = 66 ROIs that cover the entire cortex. To match this number, we simulate 24 times for each one of evenly spaced values of the global coupling parameter *G* from a suitable range, using the DMF model. We obtain 2 sets of simulations, one using the SC matrix and one using the EC matrix to set the underlying connectivity between the 66 regions. We also compare our results to a heuristically enhanced SC, SC_e_, where homotopic connections are added using a range of unit weights *D*. Each dataset is tensorized by computing dFC matrices inside of rectangular sliding windows *w* of width 2 minutes (60 frames) which are moved by one frame, resulting in *W dFC*(*w*) matrices for each subject/simulation. These matrices are concatenated into tensors of dimensions *N×N×W*. Entries of *dFC*(*w*) are calculated using correlation (positive and negative values), the absolute value of correlation (non-negative values) and mutual information (non-negative values). Each tensor is then decomposed into sets of spatial (communities) and temporal (time courses) features.

In a first step, in order to obtain community templates, we decompose the empirical tensors using different thresholds *θ*, binarizing the tensors in the case of non-zero thresholds. Since the rank of the tensors is unknown, we use different numbers *F* of features, where *F* ranges from 3 to 9. Hence, we obtain *F · S* spatial (and temporal) features for each *F*. We do not expect the temporal features to have anything in common except some general dynamic properties, so we continue with only the spatial features and cluster them, calculating the silhouette value as a quality measure for each instance of clustering. We choose the combination of *F, K* and *θ* that yields the most well-defined clusters, i.e. the highest silhouette value. The means of those clusters are the templates (i.e., *K* is the number of templates). In a second step we extract features from simulated data (*S* = 24 runs, different values of *G* and *D*) using the same *F* and *θ*, and compare each feature to the templates. We calculate an overall match for each value of *G* or *D* by taking the mean over the maximum match of each feature with any of the templates.

## 3 Results

### 3.1 A method for extracting RSNs from single subject data

Our general goal is to understand the spatio-temporal dynamics of human resting state (RS) fMRI, using time-dependent, or dynamic, FC (Chang and Glover, 2010; Hutchison et al., 2012; Allen et al., 2012; Liu et al., 2013). More specifically, we want to assess the contribution of stationary dynamics to the emergence of RSNs. To this end, we combine data analysis of long-range connectivity and and a whole-brain modelling approach (Deco et al., 2014) to investigate whether the dynamics of the model can reproduce the empirical data. We apply tensor decomposition (Cichocki et al., 2014), a method that allows us to simultaneously consider spatial and temporal structure of FC. We compute pair-wise dynamic FC (dFC) using mutual information (MI) (Kraskov et al., 2004) inside of overlapping sliding time windows (Chang and Glover, 2010; Kiviniemi et al., 2011; Hutchison et al., 2012; Allen et al., 2012; Leonardi and Van de Ville, 2013a), obtaining a dFC matrix, *dFC*(*w*), for each window *w*. These matrices are then concatenated along the temporal dimension into a 3-way-tensor (figure 2B). Additionally, we apply a binarization threshold in order to reduce noise. This is done for both empirical as well as simulated data.

By factorizing each subject’s (or simulation run’s) tensor, we find a representation of dFC in which a low number of communities superimpose linearly. Put another way, for each window *w* and *dFC*(*w*), we can determine how strongly each community contributes to the dFC-structure. This yields a time course of weights. We extract *F* = 3, 4*,…*9 communities/time courses (features) (Beckmann and Smith, 2004; Mantini et al., 2007) from each of *S* = 24 subjects which gives a pool of *F · S* communities. These are then grouped into *K* = 3, 4*,…*10 clusters using K-means clustering in order to find common patterns. Using this approach we can assess whether a true cluster structure is present, without enforcing independence between communities as in ICA, and determining under which circumstances this cluster structure emerges.

Goodness of clustering is assessed using the silhouette value (section 2.7); a high value indicates that the clusters represent the data well, i.e. cluster centers can be seen as “prototype communities”, or templates, that can be used on the whole group of subjects. In order to validate our results, we compare them against surrogate data and consider the difference in clustering performance. The surrogate data do no exhibit any long-term correlations, so any clustering structure is due to their Fourier spectrum, autocorrelations, and due to the method itself. We find that *F* = 3 and *K* = 4 results in the maximum value of 0.54. Phase randomized surrogate data (section 2.6), on the other hand, only reach 0.08. This indicates that, while there are clearly still a lot of inter-individual differences, assuming a cluster structure is supported by the data.

Figure 3A shows the difference between silhouette values for real and surrogate data for all combinations of *F* and *K*. Panel B visualizes the clusters by showing the correlations between communities for the highest difference in silhouette values. The clusters are quite well separated and correspond to four common patterns that are similar to previously described resting state networks, namely the default mode network, somatomotor network, right and left control networks, and visual network (figure 4).

**Figure 3:**
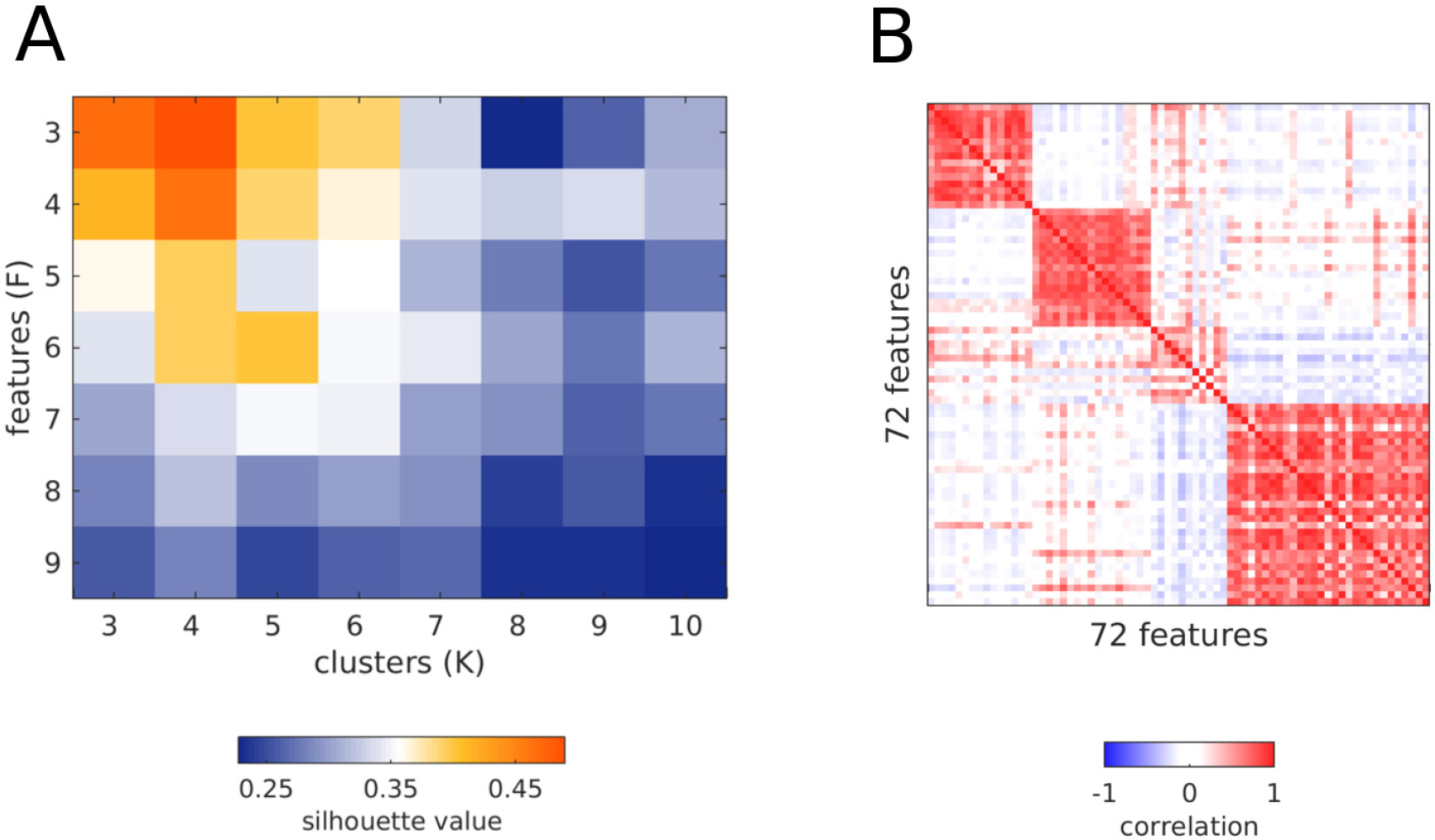
Result of template extraction with optimal parameter settings. **A** Silhouette values for clustering all features extracted from MI tensors for the best threshold, i.e. 98th percentile. **B** Correlation between the same features, ordered by assignment to *K* = 4 clusters (maximum silhouette value in **A**). The clusters are in the same order as the templates in figure 4.

**Figure 4:**
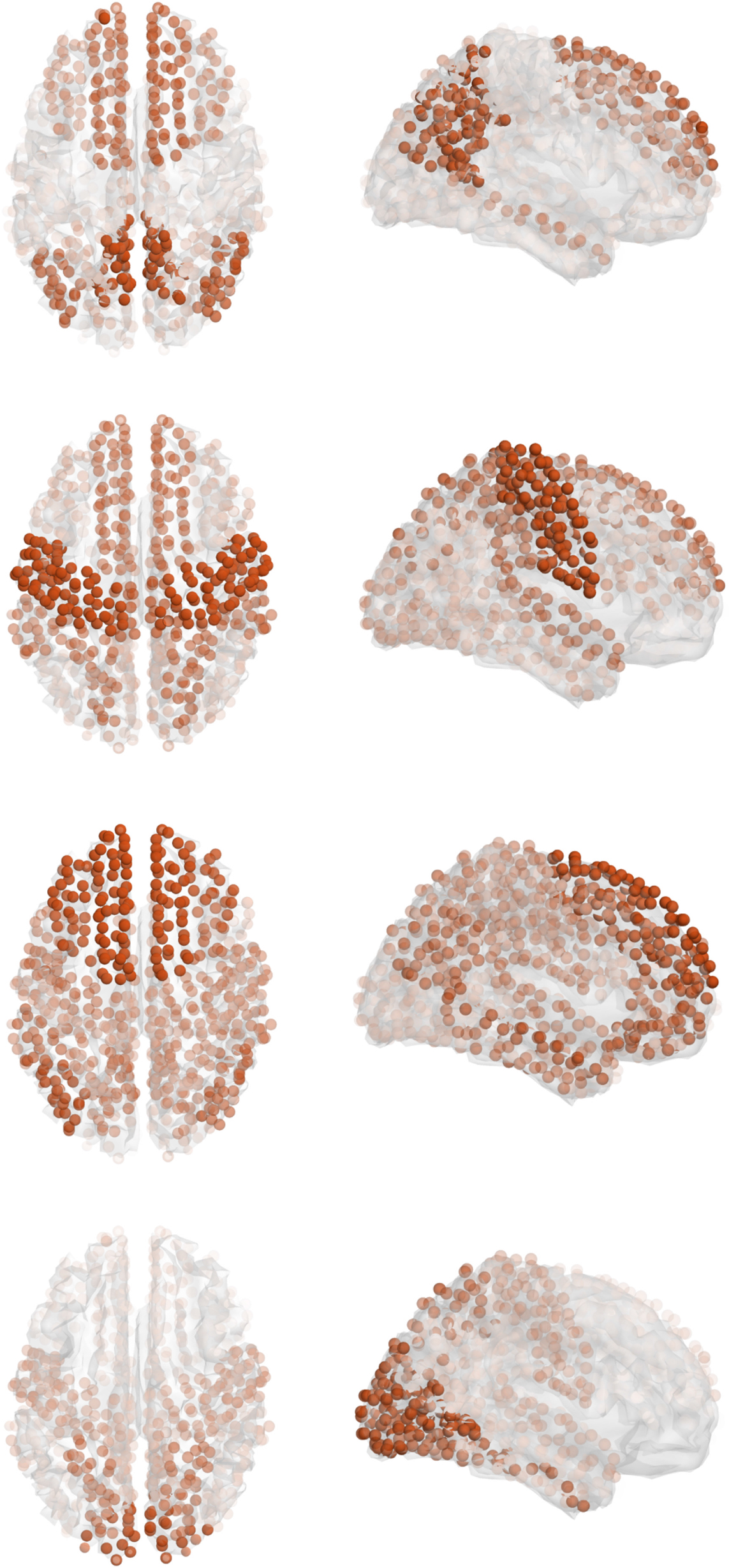
Templates plotted in 3D. From top to bottom: default mode network, somatomotor network, control networks, visual network. Although the templates are obtained at a resolution of 66 cortical regions, for better visualization we use 998 centers of mass, each of which is clearly assigned to one of the 66 areas. The opacity of each sphere is proportional to the weight assigned to it in the spatial map of the template.

Our approach is aimed at extracting patterns that explain the largest part of the data, i.e. those that occur most frequently both on the individual and the group level. We therefore wondered if our method misses those patterns that are more transient and could potentially differ greatly from the average FC, or whether they are simply not detectable at this level of resolution. We examined more closely the case where *F* > 4. Our first observation is that no additional cluster arises (see figure 3A), meaning that, if there are additional features, they do not generalize, and hence, we have no way of validating them. Our second observation is that, when examining the additional features, we find that they still resemble the four templates that we find, i.e., it happens frequently that one subject has two (or more) communities pertaining to the same cluster. We conclude that the poorer clustering performance in these cases is not so much due to inconsistent communities, but rather, because valid clusters get broken up arbitrarily. The fact that we reliably find four clusters and that the cluster centers correspond well to RSNs found in the literature speaks to the validity and robustness of our parameter selection and of this methodological approach as a whole.

### 3.2 Mutual information and a high threshold produce communities that generalize well

We use MI to compute dFC (Kraskov et al., 2004) because we find that MI is a better choice than correlation, based on clustering performance as well as reconstruction fits. Additionally, we apply a binarization threshold to the MI values, keeping only the highest ones, in order to reduce noise, which is in the following demonstrated to be necessary in order to obtain the desired generalized communities.

We start out without applying a binarization threshold, computing pairwise dFC values with Pearson correlation. Decomposing these tensors and clustering the resulting communities, we obtain the best silhouette value at *F* = *K* = 3 with a value of 0.31 for real data, and 0.19 for surrogates. This means that most of the cluster structure is due to spectral properties and autocorrelation of the data rather than real dFC. In comparison, when using MI without a threshold, we obtain the best value at *F* = *K* = 3 with 0.45 for real and 0.10 for surrogate data.

One advantage of using MI is that it allows us to apply an additional constraint when decomposing the data, i.e. non-negativity. In order to exclude that the poor result obtained with correlation is due to this, we repeat the procedure replacing the negative correlations with their absolute values. Again, *F* = *K* = 3, and we obtain 0.44 for real data, i.e. just as high as for MI, but again, 0.19 for surrogate data.

Taken together, clustering can be improved by applying the non-negativity constraint for both correlation and MI, but when using correlation, also the surrogate data exhibit more of a cluster structure, which makes MI the better choice in this application.

Another way to improve clustering is to reduce noise and thereby mitigate the effect of inter-individual differences. Furthermore, this could help remove spurious correlations introduced by spectral content below the window length (Leonardi and Van De Ville, 2015). In principle, it would be best to apply significance testing and keep only the significant dFC pairs/windows, however due to the large number of windows and pairs, this is computationally not feasible since we would have to generate thousands of surrogate tensors to reach a satisfying significance level, which would prove prohibitive in terms of computation time and storage space. Therefore, we just apply different thresholds and decompose the resulting binary tensors. We use different percentiles as thresholds, *θ* = 0, 75, 80, 90, 91, 92,…99, and compare the results, using the silhouette value.

For correlation, the absolute value of correlation, and MI, the thresholds are determined to be the 98th, the 97th and the 98th percentile, respectively. For all measures, *F* = 3 and *K* = 4. Correlation reaches a maximum mean silhouette value of 0.54 with a corresponding surrogate value of 0.13. The two non-negative measures produce better and equally good results: using the absolute value of correlation, we obtain silhouette values of 0.59 and 0.13 for real and surrogate data, respectively; for MI, the values are 0.54 and 0.08 (these templates are the ones shown in figure 4).

Apart from the clustering performance, we consider the reconstruction fits that quantify how well the extracted features describe the original tensor. At their best thresholds, MI and absolute value of correlation reach average fits of 0.39 and 0.48, respectively. The corresponding surrogate tensors yielded 0.09 and 0.34, confirming that while using the absolute value of correlation results in a good decomposition performance, the surrogate tensors constructed in this way also exhibit a lot of structure. Taken together, MI is the better choice.

### 3.3 Dynamic mean field model reproduces RSNs

In the next step, we use the templates shown in figure 4 to determine whether the dynamic mean field (DMF) model (figure 1) can explain the emergence of clearly discernable RSNs, in the sense that they exhibit their own time courses to a degree sufficient for the tensor decomposition algorithm to pick them up. We run *S* = 24 simulations of the same length as the empirical data and use the parameters previously determined with the empirical data to the resulting tensors, i.e. *F* = 3 and *θ* = 98. The only free parameter is the global coupling, *G* which is a factor by which the connectivity matrix is multiplied and is hence related to the overall amount of activation in the system. We determine for each of the *F · S* = 72 extracted features the maximum correspondence to any of the *K* = 4 empirical templates by computing Cohen’s Kappa (section 2.8). We then use the mean across all features and all runs as a measure for the match between features (simulated) and templates (empirical) and thus of how well the model reproduces the empirical data. As before, we consider the difference to matches obtained using surrogates, i.e. of features extracted from tensors that are calculated from phase randomized simulated data to empirical templates.

We use the parameters obtained from the empirical data instead of running the same parameter selection procedure on the simulated data. This is because the simulations run on an average connectivity matrix and therefore, clustering across simulation runs does not make sense; the variance in FC structure is small across runs for a given value of *G*.

Figure 5A shows the overall overlap between simulated features and templates depending on the global coupling parameter. The 95% confidence intervals of real and surrogate data overlap for most values of *G*, except for *G* = 1.9, 2.5 − 2.7, 2.9, and 3.1 − 3.7, the latter values matching the region in which the fit of the average FC is best. For these values, the simulated data can be shown to contain FC patterns that match empirical data to a higher degree than surrogate features. However, even the best match at *G* = 3.7 is moderate. We find that the main problem is that the features are not symmetrical, but rather, lateralized, which is in line with results by Messé et al. (2014). For example, all features that match with the somatomotor template have their members in the right hemisphere. Surprisingly, the visual network, which has proven to be the most clearly pronounced one in the empirical data, is not found at all (see figure S1).

**Figure 5:**
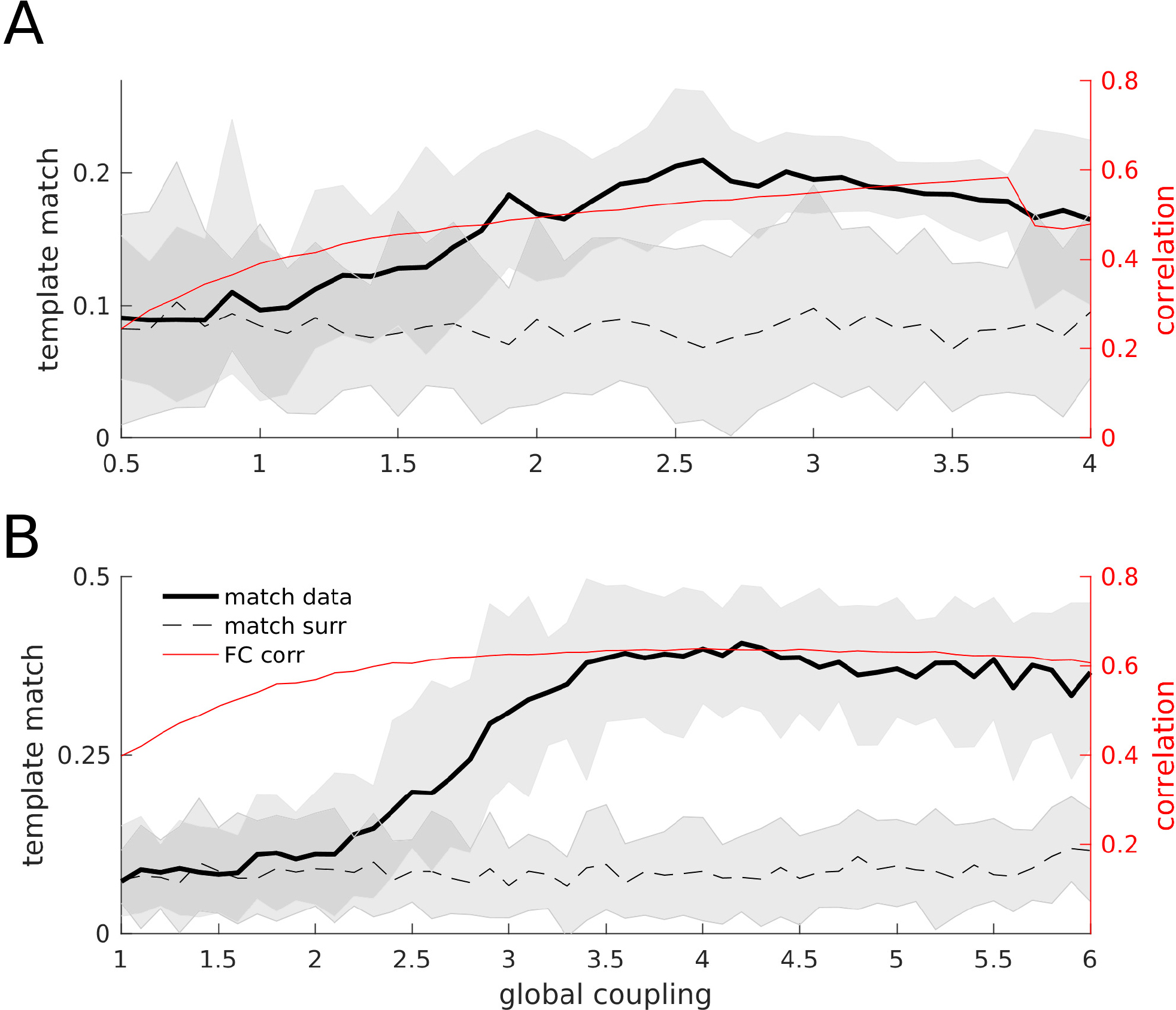
Overall template matches in comparison to correlation fits between average FC matrices, results for simulations using SC (panel **A**) and EC (panel **B**). Note the different scales on the left y-axes. Black curves: template matches (left axes), thick black curve is real data, dashed line surrogate data. Shaded areas indicate the 95% confidence interval. Red curves: correlation of average FC matrices (right axes).

SC is derived by applying fiber-tracking algorithms to diffusion tensors obtained via dwMRI. We use the method developed by Gilson et al. (2016), briefly described in section 2.9, to obtain from this SC an effective connectivity (EC) for our dataset. This EC contains meaningful weights as well as directionality information. Importantly, only the weights that are also present in the SC are tuned by the procedure, plus the weights on the secondary diagonal to account for homotopic connections that are not represented well by fiber tracking.

Figure 5B shows the overall overlap between simulated features and templates when EC is used (note that the scale on the y-axis is different). The non-overlapping region of the 95% confidence intervals of real and surrogate data covers a wider range of *G* (2.9 to 6.0), the maximum being at *G* = 4.2. Figure 6 shows the features extracted at this point next to the templates in vector form. They are symmetrical and agree well with the templates although the overlap in figure 5B may still seem somewhat modest.

**Figure 6:**
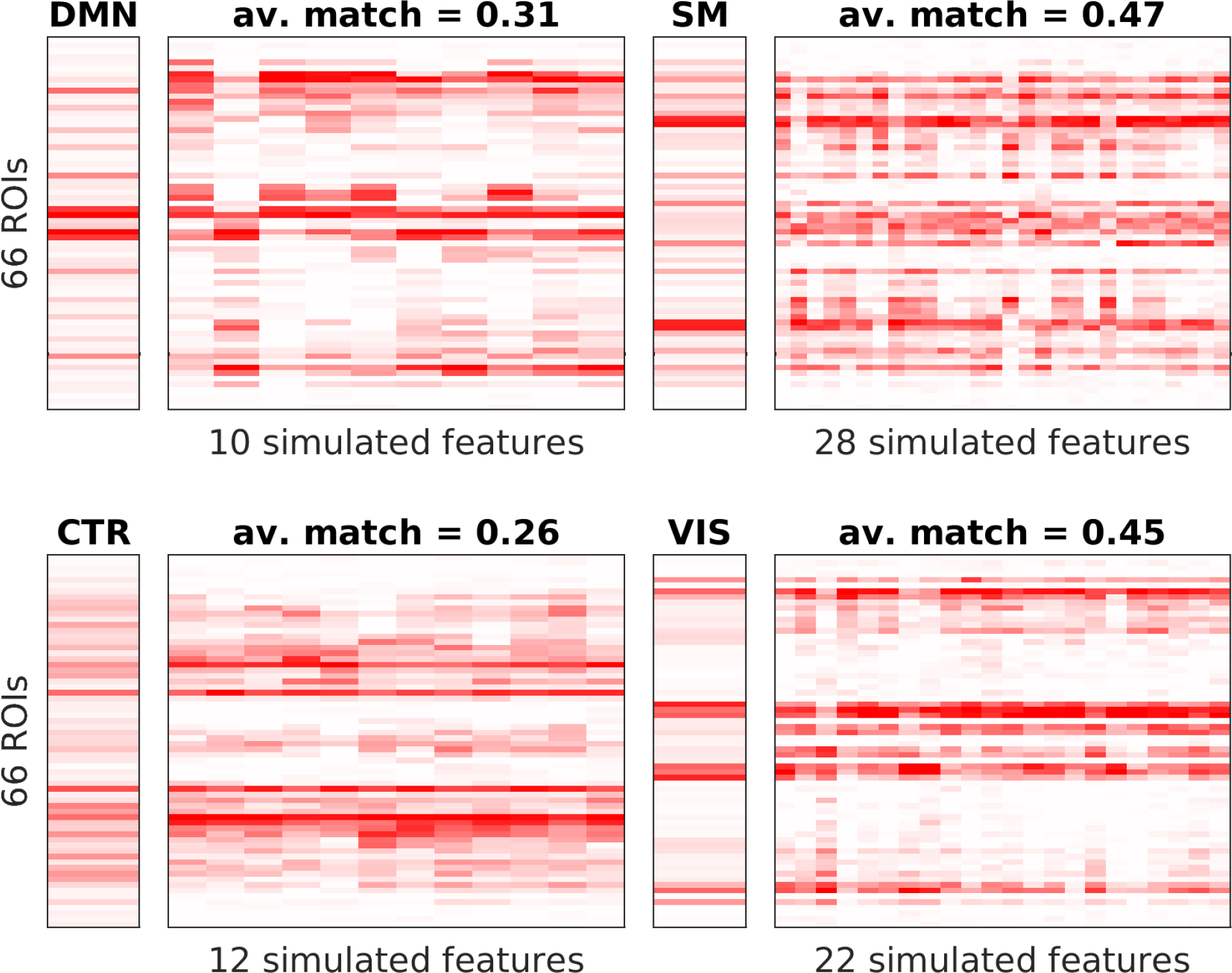
Spatial features extracted from 24 simulation runs using EC as underlying connectivity and the *G* with the best match (figure 5: *G* = 4.2) with *θ* = 98th percentile and *F* = 3. Again, each simulated spatial feature is plotted next to the template with which it exhibits the higest match according to the *κ*-value (see section 2.8). Templates correspond to DMN - default mode network, SM - somatomotor network, CTR - left and right control networks, VIS - visual network. For each set of features, the number of matched features and the average overlap for only this set is given.

We explain this better match by considering panels B and C of figure 7. Several differences between SC and EC are immediately obvious, although we first note that both of them differ greatly from the FC shown in panel A; the correlation values between SC and FC as well as SC and EC are both 0.35, speaking to the strong network effect exerted by the DMF model simulations. First of all, homotopic connections that are missing in SC are prominent in EC. This can explain the symmetry in the features obtained from EC-based simulations. Following Messé et al. (2014), we test how the fit (both to average FC and RSNs) can be improved just by adding these connections, obtaining a heuristically enhanced SC, termed SC_e_. We test 50 different weights *D* for these connections, covering a wide range of values present in the SC (see section 2.9). We find that the fit to the average FC is improved to the level that EC achieves (SC: 0.58, EC: 0.64, SC_e_: 0.65), consistent with Messé et al. (2014). This fit is similar for all tested values of *D* (see figure S2). We find that the match between simulated features and templates reaches a maximum value that lies between that for SC (0.21) and EC (0.41), i.e. 0.32. This means that with a very simple adjustment, we could achieve a substantial improvement.

**Figure 7:**
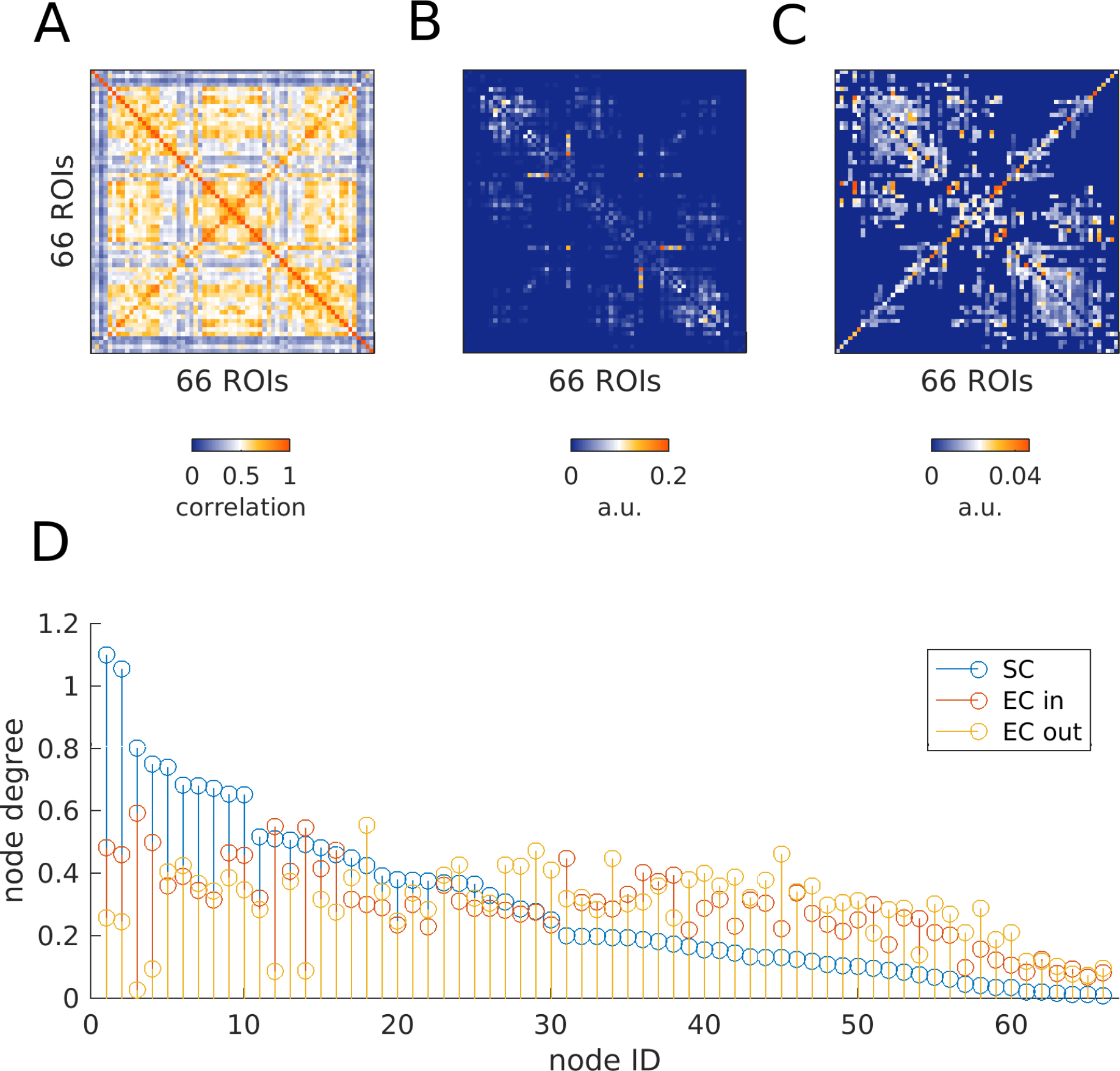
Connectivity matrices. **A** Functional connectivity matrix averaged over 24 subjects. Each entry is the correlation over the entire time course of a pair of ROIs. **B** Average structural connectivity matrix from the same 24 subjects. **C** Effective connectivity matrix derived from the FC, using the method in section 2.9. **D** Degrees of the nodes. For SC (blue), in and out degree are identical. Here, nodes are ordered according to their degree. EC nodes (in-degree: orange, out-degree: yellow) are shown in the same order as for SC.

However, while the resulting features are now symmetrical, there are still no communities that match the prominent visual RSN (figure S3). Furthermore, the condifence intervals of overlaps obtained from real vs. surrogate data overlap for most values of *D* (figure S2), as was the case for SC.

Furthermore, the weights are more uniformly distributed in EC, making it appear more dense. Panel D of figure 7 reveals that the node degrees are largely equalized, such that most of them range in the middle of the distribution. This explains the different ranges of *G* that can be used in the simulations. For SC, the nodes with the largest degree cause the firing rates in the corresponding excitatory pools to rise to the point where the inhibitory pools cannot compensate for them any longer, and the asynchronous, low-activity regime of the system becomes unstable. The more uniform node degree distribution in the EC matrix allows for higher values of *G* and thus, the communities become more pronounced due to improved signal to noise-ratio. While the difference between the highest and lowest node degrees is smaller in SC_e_ than in SC, we did not observe the same kind stabilizing effect as with EC, although the increase in maximum mean firing rate was less extreme.

The EC matrix is also non-symmetric. This property likely further contributes to the stability of the simulations and more generally to the more realistic shape of extracted communities, both on the spatial and temporal level because it allows for a more diverse propagation of the activity through the entire network.

## 4 Discussion

Our goal was to characterize recurring patterns of FC extracted from human resting state (RS) fMRI and to determine whether a noise-driven stationary mean field model (Wong and Wang, 2006; Deco et al., 2014) can reproduce them. Using tensor decomposition (Cichocki et al., 2009), we identify four communities that generalize across subjects and resemble known RSNs (Fox et al., 2005; Beckmann et al., 2005; Mantini et al., 2007). We utilize temporal information by computing pairwise dynamic FC (dFC) in sliding windows of 2 minutes to build our tensors. We compare three dFC measures: correlation, absolute value of correlation, and mutual information (MI). We determine the dFC measure, number of extracted features *F*, number of templates *K*, and binarization threshold *θ*, that yield the best clustering performance, and take cluster centers as templates. We find that using a low number of features (*F* = 3) and of clusters (*K* = 4) combined with a high threshold *θ* = 98th percentile) applied to dFC calculated from MI works best (figures 3, 4).

We determine the range of global coupling for which the match of communities extracted from the model data to the templates is maximal (figure 5). We compare two underlying connectivities of the model: dwMRI derived structural connectivity (SC) and model-based effective connectivity (EC), which estimates directionality and weights of connections using time-shifted covariances (Gilson et al., 2016). EC produces more realistic FC patterns than SC alone, which is not explained by the homotopic connections. Particularly, the RSN corresponding to the visual system is the strongest component in the empirical data, but is not found in the simulated data unless EC is used.

### 4.1 Few data points are sufficient to recover RSNs

We find that applying a threshold and binarizing the tensors is crucial in order to obtain a decent clustering performance. The best threshold of *θ* = 98 translates to using only the 2% biggest dFC values in the decomposition and they are sufficient to extract RSNs (figures 3 and 4).

This is in line with Tagliazucchi et al. (2012, 2016), who transformed RS fMRI time courses into a point process, reducing the data by 94% and keeping only extreme events. Also there, the authors were able to recover RSNs. They suggest that “avalanching events which involve short and long range cortical co-activations” explained their results. A related approach is described in Karahanoğlu and Van De Ville (2015), where the authors model fluctuations in the fMRI signal as a consequence of underlying events (“innovation signals”). The fact that these results are so similar suggests that periods of highly structured FC (many big dFC values) are equivalent to periods of high variance in the BOLD signal (many ROIs crossing the threshold set for the point process). Indeed, figure 8 shows that the time courses of RSNs fluctuate strongly, the peaks occurring in windows with many supra-threshold dFC pairs. It is not immediately evident to what degree RSNs overlap in time, and this deserves further attention in future studies, but it seems clear that they overlap at least to some, not negligible degree. We conclude that the peaks of these fluctuations represent periods of high modularity which are detected by the factorization algorithm. Modularity and related graph metrics like efficiency or path length, have previously been shown to dynamically fluctuate (Tagliazucchi et al., 2012; Zalesky et al., 2014; Betzel et al., 2016a).

**Figure 8:**
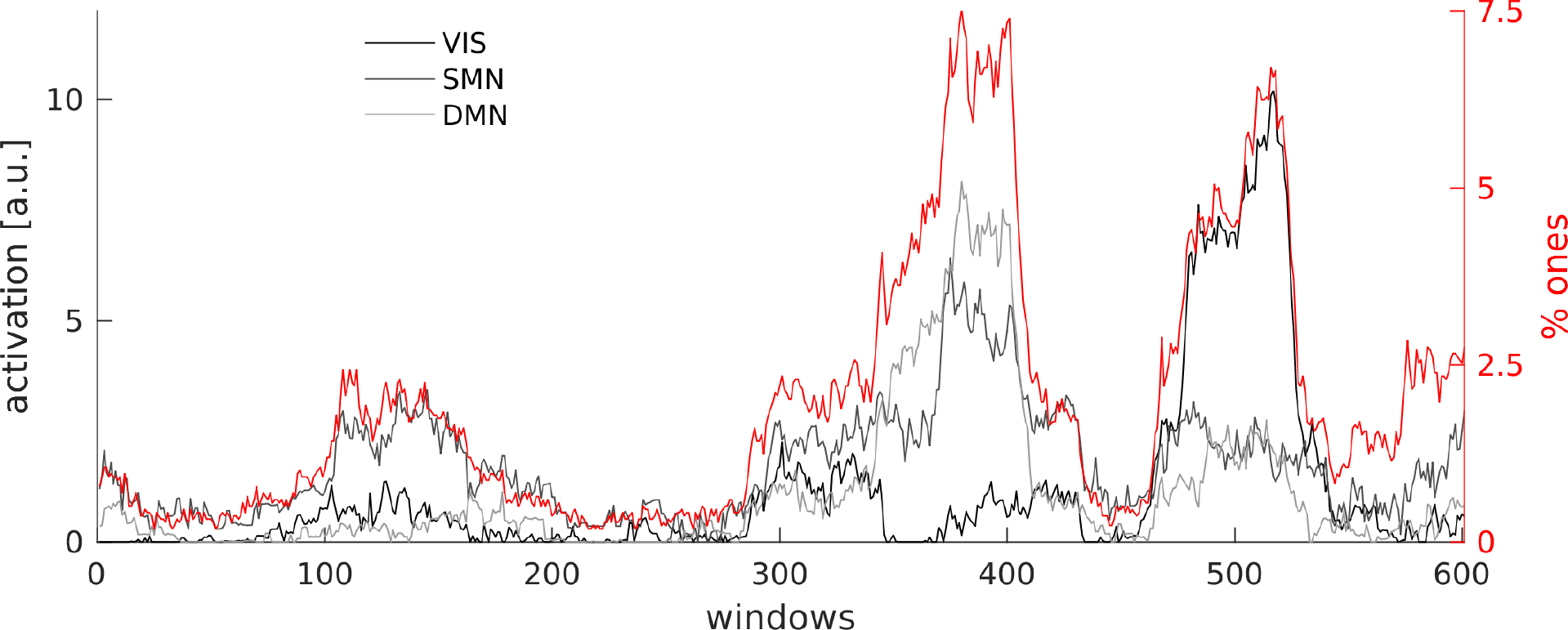
Time courses of the RSNs from one subject extracted via tensor decomposition (using MI and *θ* = 98th percentile) together with the percentage of supra-threshold dFC values in the window (ones in he binary tensor’s slices). The RNSs are identified as VIS-visual, SMN-somatomotor, DMN-default mode network.

A complimentary approach to modelling dFC as arising from an underlying point process is provided by Mitra et al. (2014). The authors explain observed data in terms of waves of activity that propagate from regions acting as sources to others acting as sinks, involving the entire cortex. This results in stereotypical dFC patterns derived from average FC. Our results show that short time scales at the level of the point processes and also the EC method used here contain the same information as long time scales at the level of sliding windows and the entire scan duration in the sense that RSNs can be recovered in all cases. In the simulated data, this property is reproduced by virtue of an adequate underlying connectivity (the EC) and a rather weak hypothesis on the dynamics, since the DMF model is stationary.

From this point of view, the fact that we can extract RSNs from a low number of windows is merely a consequence of fluctuations in the variance/signal to noise-ratio in the underlying signal; in other words, we could extract RSNs from *any* set of windows (provided it is sufficiently large).

Furthermore, Messé et al. (2014) showed that a stationary model assuming fluctuations around a single fixed point, i.e., the average FC, predicts spatial structure of FC more accurately than more complex models, even when dFC is included. This is in parallel to what we show here: even a model that possesses only one attractor can produce time-varying FC patterns that are picked up by a decomposition algorithm.

More explicitly, this goes against the notion that the observed infraslow modulations are a signature of switching between distinct dFC patterns as proposed for example in Hansen et al. (2014). There, the authors correlate all dFC matrices occurring over time, yielding a recurrence plot which shows a checkerboard-like structure: long lasting periods of highly autocorrelated dFC patterns are followed by a sharp drop in correlation of subsequent dFC matrices. The time scale matches that shown in figure 8. This is taken to correspond to different states, perhaps RSNs or distinct combinations thereof. Our results suggest that on this time scale, there are only two “states” which exist at the two ends of a continuum: windows with high vs. low modularity.

### 4.2 Why we use tensor decomposition

In this study, we use a method that is relatively unknown in neuroscience, i.e. tensor decomposition (Beckmann and Smith, 2005; Cichocki, 2013; Leonardi and Van de Ville, 2013a; Leonardi and Van De Ville, 2013b; Gauvin et al., 2014; Ponce-Alvarez et al., 2015). There are several reasons for our choice. First of all, with a resolution of 661 time frames and 66 ROIs, ICA cannot be readily applied. We would have to use temporal ICA (Calhoun et al., 2001) instead of the more widely used spatial ICA (McKeown et al., 1998; Beckmann and Smith, 2004; Calhoun et al., 2009), making it difficult to compare to studies that identify communities (Beckmann et al., 2005; Damoiseaux et al., 2006; De Luca et al., 2006; Mantini et al., 2007; Van den Heuvel and Hulshoff Pol, 2010). However, the low spatial resolution was necessary in order to assess whether our specific model - the dynamic mean field model (Deco et al., 2014) - does indeed have the capacity to reproduce RSNs. Apart from that, investigating specifically long-range connections in a whole-brain approach has its own merit. In any case, tensor decomposition can be seen as a generalization of PCA or SVD and is thus a fairly generic data analysis technique.

Our second reason is that this method allows us to use temporal fluctuations of FC explicitly, but without making any assumptions. Indeed, non-negativity is the only constraint we apply, and it is natural to do so when using a non-negative FC measure like MI. In other words, while ICA starts with the BOLD time courses, we use pair-wise FC, investigating time-evolving community structure explicitly. The result is that we get spatial and temporal features at the same time, without having to assume independence in either dimension. Previous publications have made use of dFC explicitly by applying higher-order SVD (HOSVD) to their tensors (Leonardi and Van de Ville, 2013a; Leonardi and Van De Ville, 2013b). However, in contrast to CPD as used here, a clear correspondence between spatial and temporal features is not easily established in HOSVD, because, as in SVD for matrices, the resulting subspaces are required to be orthogonal to each other. This leads to spatial features being interpreted as connectivities (“eigenconnectivities”) and not as communities of ROIs.

Third of all, and importantly, this method allows us to extract features from single subjects or simulations, making it unnecessary to map group results back to the individual like with dual regression in group ICA. It is straightforward to match extracted features to a group template and estimate how well a subject agrees with the group average, which is summarized in the silhouette value used here to validate clustering results and tune parameters. This also provides us with a mechanism to select the decomposition parameters (*K, F, θ*) and guarantess robustness of the results in a data-driven manner. A similar approach (with hierarchical clustering) was used by Ponce-Alvarez et al. (2015). Alternatively, the third dimension can be used for the subject dimension, as done in Leonardi and Van de Ville (2013a) and typical for tensor probabilistic ICA (Beckmann and Smith, 2005). However, there, dFC matrices are again flattened into vectors, leading to “eigenconnectivities” as mentioned above. This is also the approach taken in Leonardi et al. (2013), where PCA was applied in order to extract dFC patterns, but here, the resulting matrices were additionally concatenated in the subject domain, reducing the problem to two dimensions. Thus, K-means clustering is an alternative approach to dealing with the subject dimension. In principle, it would be conceivable to use 4-way-tensors, but the feasibility and merit of such an approach remains to be explored.

### 4.3 Correlation versus mutual information

Another perhaps slightly unusual choice is to favor MI over correlation as a measure of FC. It has, however, several advantages. First, it is a non-negative measure which allows us to further constrain the tensor decomposition, leading to more reliable results. Second, spurious dFC structure in correlation-based tensors is reduced, as consistently shown by our results.

The reason for this difference has to lie in the spectral properties of the data, because they are preserved in the surrogate data. We found many time windows that contain outliers, i.e. a handful of time points with much higher activity than the rest. MI as a non-parametric measure and as computed here (Kraskov et al., 2004) is robust against these outliers, but correlation assumes normality and therefore overestimates dFC in such windows. We hypothesize that these outliers are a result of global slow fluctuations in the signal that are preserved in the surrogates. Another factor which could contribute to this difference in performance are possible spurious correlations introduced by spectral content below the window length (Leonardi and Van De Ville, 2015), the effects of which MI might help to mitigate.

This observation calls into question the usefulness of Pearson correlation for investigating dFC despite its popularity (Allen et al., 2012; Hutchison et al., 2013; Hansen et al., 2014). Concerns have been addressed in some publications (Lindquist et al., 2014; Hindriks et al., 2015), and cross-validation with appropriate surrogate data is highly commendable.

One feature that a non-negative measure might seem unable to address is that of anti-correlation between DMN and task-positive network, a hallmark of RS fMRI (Raichle et al., 2001). However, the time scale examined here is not the same as that of anti-correlations, leading to a view where they are embedded in a global, slower modulation. Furthermore, anti-correlated activation could in principle still show up in the MI time courses.

### 4.4 Using state-of-the-art connectivity matrices

In SC, each connection is symmetric and its weight is determined by the number of streamlines detected by the tracking algorithm. However, it is known that many long-range connections are missed by these algorithms because of crossing fibers, notably in the region of the corpus callosum but also connections between frontal and occipital regions. Therefore, a lot of the interhemispheric connections between homotopic areas are missing. Furthermore, the results of fiber tracking do *not* allow direct inference of the weight of a connection. Lastly, fiber tracking results exhibit a high variability across subjects and sessions (Jones et al., 2013; Jeurissen et al., 2013). Therefore, DTI matrices should not be read as reflecting true connectivity, but rather, the weights reflect how easy it is for the fiber tracking algorithm to establish a streamline between two locations, given the diffusion data and the assumptions incorporated in the algorithm. For example, here we use matrices that were obtained using sophisticated error correction procedures, like hypotheses about the relationship between size of the grey-white-matter-interface and the space that a fiber can occupy for any given region (Schirner et al., 2015).

On top of these shortcomings, we know that, *in vivo*, the asymmetry of the underlying connectivity shapes the dynamics of the system. To estimate weights and directionality, we need suitable observables and a dynamical model. We use the conceptual framework of EC (Friston, 1994), which describes the influence that one cortical area has over another. It depends, in principle, on synaptic plasticity, previous brain activity, and neuromodulation. Therefore, *in vivo*, EC changes dynamically depending on the experimental condition, individual differences, etc.

The method developed in Gilson et al. (2016) allows us to extract a single (group level) EC from our data that remains unchanged for all simulations. Directionality and weights of connections are estimated using time-shifted, i.e., asymmetric, covariances as observables, as opposed to correlations.

Many other studies have dealt with the estimation of EC, but this approach has several hallmarks which are advantageous and interesting for this study. First of all, it can easily handle the number of brain regions used here, as opposed to DCM (Friston et al., 2014), which, moreover, is appropriate in a situation where we have a few hypothetical network configurations which we wish to compare. In the same vein, the method summarized here is among the relatively few which consider the network as a whole (Smith et al., 2011). Furthermore, strong FC can exist between nodes which are not physically connected in the SC, i.e. a strong network effect exists, as was the case for example in Deco et al. (2013) and Robinson et al. (2014). However, in those studies, the directionality of the connections was not estimated. Here, the usage of BOLD covariances with non-zero time shifts as an observable enables us to do just that, yielding an asymmetric EC matrix.

The use of EC leads to more realistic communities that are impossible to obtain with SC alone. An improved match between empirical and simulated is expected when tuning more parameters, however, the relationship between node connectivity in the DMF model (SC or EC) and the resulting FC of the data is not straightforward, as is evident from the relatively low correlation between the two; notably, this correlation is the same for SC as for EC (0.35 in both cases). One of the main benefits of using EC is that homotopic connections are strengthened, enabling realistic symmetric communities. Messé et al. (2014) found that just adding homotopic connections by setting them to a fixed value greatly improved predictive power for the average FC. We found that this result extends to RSNs extracted from dFC. We would like to make two points about this result. First of all, this approach is completely heuristic and thus, from a certain point of view, it is preferable to include a dynamical model which will give us insight about the relationship between SC/EC and FC. Furthermore, while obtaining the EC only takes a few minutes, assessing the effect of different weigths in the homotopic connections requires many hours of simulations.

One question of this study is: “To what extend are our observations explained by stationary dynamics?” Thus, testing how far we can improve our match between interesting aspects of the empirical data - here, RSNs obtained from dFC - by optimizing the underlying connectivity *without adding any nonstationarities* has its own merit. In Messé et al. (2014), where the authors asked a similar question, it was found that even when including dFC, the stationary model was still the one with the biggest predictive power. It would be interesting to see how model performances play out when using EC, because the shortcomings of SC are likely to impair modelling results very strongly. Perhaps, once this limitation is mitigated, we would favor a more complex model, as was the authors’ initial hypothesis.

We emphasize that the communities themselves do not contribute in any way to the EC optimization procedure. They are extracted using temporal information in the shape of *dFC*(*w*) matrices that are obtained on time scales that are far greater than those used in the optimization procedure. Thus, the two methodologies make completely different use of dFC. Furthermore, the template extraction which drives the adjustment of parameter setting (choice of *F, K*, dFC measure, and *θ*) is done on the single subject basis and is independent of any simulations. Nevertheless, we do believe that temporal dynamics of spontaneous BOLD activity takes place on multiple time scales. The fact that we find that the emergence of RSNs can in principle be explained by stationary dynamics does not imply stationarity on the time scale used by EC. Faster, nonstationary processes could be embedded in infraslow modulations on the level of the average FC.

### 4.5 Conclusion

We have shown that a dynamic mean field model with a single attractor that is explored through noise is sufficient to explain a lot of the spatio-temporal structure found in large-scale resting state fMRI. Noise-driven fluctuations around the average functional connectivity structure are shaped by the underlying connectivity and the simple dynamics of the model in such a way that known functional networks emerge from the temporal dynamics. This adds to previous findings according to which the average FC is reproduced by the model by demonstrating that RSNs can be extracted from the temporal dynamics of the simulated data.

These results are at odds with an understanding where the brain at rest switches in a nonstationary manner between RSNs because our model is certainly not capable of this behavior. In the simulated data, “states”, i.e. dFC matrices computed in a time window as a proxy for instantaneous FC, are not equivalent to RSNs but are always a noisy version of the average FC; still, RSNs can be extracted as common building blocks of dFC across subjects, in our case, by decomposing tensors, or “stacks” of time-dependent dFC matrices. Of course we are not claiming that the brain is stationary even during rest. Switching likely still occurs at a faster time scale and finer spatial resolution. Even more, the modulations in average FC might be embedding other processes that could be discovered when discounting this global component.

Apart from that, even this “global component” might still be meaningful. A possible interpretiation is provided by recent work on network controllability (Gu et al., 2015; Betzel et al., 2016b) which shows that a node’s topological role (node degree, centrality, etc.) strongly determines how it contributes to transitions from an initial towards a specific target state. Controllability, or the ability of a brain region to steer the network to a certain state, depends, among other things, on the underlying connectivity (SC, SC_e_, or EC in this work) and on the “passive” dynamics, i.e. the path through state space that the system would take if no controlling inputs were applied - in other words, spontaneous dynamics as investigated here. This is in line with the idea that during RS, the brain prepares for potential incoming tasks and stimuli. Using the control theory framework, we could say that the brain network and node dynamics are set up in such a way that important target states (e.g., when responding to a threat) can be reached with minimal energy cost.

Future studies should try to further characterize the temporal structure, for example determining how separate in time different networks are. This would enable us to pinpoint differences between experimental groups (different ages, gender, patient groups) and regarding tasks (e.g. attention, decision-making, motor) by describing and modelling how large-scale networks interact and how these characteristics are related to performance or clinical markers.

## Acknowledgements

The authors would like to thank Jochen Braun, Romain Brasselet, Rikkert Hindriks, Raphael Kaplan, and Gorka Zamora for valuable discussions about the work in this manuscript.

KG: Special thanks to Professor Andrzej Cichocki, Dr Zhou Guoxu, Dr Qibin Zhao, and Dr Anh Huy Phan from RIKEN BSI, Wako-shi, Saitama, Japan, for their support and supervision during my training stay in the Advanced Brain Signal Processing group in Summer 2014 during the RIKEN BSI summer program.

## Notes

Funding (authors’ initials given after grant numbers): This work was supported by the European Union, FP7 Marie Curie ITN “INDIREA” (Grant N. 606901; KG), FP7 FET ICT Flagship Human Brain Project (Grant N. 604102; MG), ERC Advanced Human Brain Project (Grant N. 604102; GD), Horizon2020 ERC Consolidator (Grant N. …; PR); the Spanish Ministry for Economy, Industry and Competitiveness (MINECO) project “PIRE-PICCS” (Grant N. PCIN-2015-079), SEMAINE ERA-Net NEURON Project (Grant N. PCIN2013-026; APA), nd ICoBAM (Grant N. PSI2013-42091-P; GD); the James S. McDonnell Foundation (Brain Network Recovery Group, Grant N. JSMF22002082; PR); the German Ministry of Education and Research (Grant N. 01GQ1504A and 01GQ0971-5; PR); the Max-Planck Society (Minerva Program; PR)

